# Genetic pleiotropy between mood disorders, metabolic, and endocrine traits in a multigenerational pedigree

**DOI:** 10.1101/196055

**Authors:** Rachel L. Kember, Liping Hou, Xiao Ji, Lars H. Andersen, Arpita Ghorai, Lisa N. Estrella, Laura Almasy, Francis J. McMahon, Christopher Brown, Maja Bućan

**Affiliations:** Department of Genetics, Perelman School of Medicine, University of Pennsylvania, Philadelphia, PA 19104; Human Genetics Branch, National Institute of Mental Health Intramural Research Program, Nation Institutes of Health, Bethesda, MD 20982; Genomics and Computational Biology Program, Perelman School of Medicine, University of Pennsylvania, Philadelphia, PA 19104; Lancaster General Health/Penn Medicine, University of Pennsylvanai Health System, Lancaster, PA 17602; Department of Biomedical and Health Informatics, Children’s Hospital of Philadelphia, Philadelphia, PA 19104; Department of Psychiatry, Perelman School of Medicine, University of Pennsylvania, Philadelphia, PA 19104

**Author notes:** Corresponding author: Maja Bućan.

## Abstract

Bipolar disorder (BD) is a mental disorder characterized by alternating periods of depression and mania. Individuals with BD have higher levels of early mortality than the general population, and a substantial proportion of this may be due to increased risk for comorbid diseases. Recent evidence suggests that pleiotropy, either in the form of a single risk-allele or the combination of multiple loci genome-wide, may underlie medical comorbidity between traits and diseases. To identify the molecular events that underlie BD and related medical comorbidities, we generated imputed whole genome sequence (WGS) data using a population specific reference panel, for an extended multigenerational Old Order Amish pedigree (400 family members) segregating BD and related disorders. First, we investigated all putative disease-causing variants at known Mendelian disease loci present in this pedigree. Second, we performed genomic profiling using polygenic risk scores to establish each individual's risk for several complex diseases. To explore the contribution of disease genes to BD we performed gene-based and variant-based association tests for BD, and found that Mendelian disease genes are enriched in the top results from both tests (OR=20.3, p=1×10^−3^; OR=2.2, p=1×10^−2^). We next identified a set of Mendelian variants that co-occur in individuals with BD more frequently than their unaffected family members, including the R3527Q mutation in *APOB* associated with hypercholesterolemia. Using polygenic risk scores, we demonstrated that BD individuals from this pedigree were enriched for the same common risk-alleles for BD as in the general population (β=0.416, p=6×10^−4^). Furthermore, in the extended Amish family we find evidence for a common genetic etiology between BD and clinical autoimmune thyroid disease (p=1×10^−4^), diabetes (p=1×10^−3^), and lipid traits such as triglyceride levels (p=3×10^−4^). We identify genomic regions that contribute to the differences between BD individuals and unaffected family members by calculating local genetic risk for independent LD blocks. Our findings provide evidence for the extensive genetic pleiotropy that can drive epidemiological findings of comorbidities between diseases and other complex traits. Identifying such patterns may enable the subtyping of complex diseases and facilitate our understanding of the genetic mechanisms underlying phenotypic heterogeneity.

## Introduction

Psychiatric disorders frequently co-occur with other medical illnesses, markedly reducing patients’ quality of life. Individuals with mood disorders have higher levels of early mortality than the general population, in part due to comorbid medical disease (Parks et al., 2006). In addition, individuals with a higher burden of medical illness display not only a higher occurrence but also an increased severity of psychiatric symptoms (Beyer et al., 2005). Bipolar disorder (BD) is a highly heritable mood disorder characterized by recurrent periods of depression and mania. Individuals with BD have increased rates of asthma, diabetes, hyperlipidemia, epilepsy and thyroid disease, among other diseases (Forty et al., 2014).

Historically, increased rates of medical illness in patients with psychiatric disorders had been attributed to the side effects of antipsychotic medication or to a reduced ability to maintain a healthy lifestyle. However, recent evidence suggests that shared genetic risk loci or common biological pathways may underlie the pervasive pleiotropy between psychiatric and non-psychiatric disorders (Bulik-Sullivan et al., 2015; Pickrell et al., 2016; Prieto et al., 2016; Sivakumaran et al., 2011). Pleiotropy can be identified at the level of individual alleles, or genetic correlations between disorders can be calculated genome-wide to quantify the proportion of shared associated loci between traits (Bulik-Sullivan et al., 2015).

Comorbidity arising from pleiotropic loci has been noted in Mendelian disorders, with a significant number of Mendelian disease-causing variants leading to complex phenotypes (Zhu et al., 2014). In individual-level data gathered from medical records of over 110 million patients, Mendelian variants were found to contribute non-additively to risk for a subset of complex diseases (Blair et al., 2013). Furthermore, common variants associated with complex disease are enriched in Mendelian disease genes (Blair et al., 2013). Shared genetic influences between common complex traits have also been identified. Using data from genome-wide association studies (GWAS), several groups have identified loci underlying multiple traits (Pickrell et al., 2016; Sivakumaran et al., 2011). Such cross-phenotype associations (Solovieff et al., 2013) have been found even between distinct traits; for example, a nonsynonymous variant in *SLC39A8* is associated with both schizophrenia and height, among others (Pickrell et al., 2016).

Genetic correlations have demonstrated the shared genetic influences between multiple clusters of diseases, including significant correlations between psychiatric disorders (Gale et al., 2016; Hammerschlag et al., 2017; Lee et al., 2013; Lo et al., 2017). Shared genetic etiology has been identified between BD and both schizophrenia and major depressive disorder (Lee et al., 2013), suggesting extensive biological pleiotropy between these psychiatric conditions (O'Donovan and Owen, 2016). Given the polygenic nature of these disorders (Wray et al., 2014), this result is unsurprising, as polygenicity is consistent with comorbidity and pleiotropy (Gratten et al., 2014). Gratten et al. (2014) also notes that genetic correlations between BD data sets are more variable, possibly suggesting greater genetic heterogeneity within BD compared to other psychiatric phenotypes.

Population isolates are frequently utilized in genetic studies of disease in order to reduce the genetic and phenotypic heterogeneity found in outbred populations. Disease-gene identification in population isolates usually attempts to identify both common and low-frequency variants, observed within and across families. Moreover, founder effects can lead to an increase in allele frequencies for many deleterious alleles and clusters of deleterious variants on shared haplotypes. Genetic studies of the Old Order Amish led to the identification of over 200 Mendelian disease loci (Puffenberger et al., 2012; Strauss and Puffenberger, 2009; Strauss et al., 2012), and the same population has been extensively utilized in studies of complex metabolic and psychiatric diseases (Georgi et al., 2014; Kember et al., 2015; Rampersaud et al., 2007; Steinle et al., 2005; Strauss et al., 2014; Xu et al., 2017).

The Amish Study of Major Affective Disorder (ASMAD) is a large extended pedigree collected over 30 years ago with an initial focus on bipolar disorder. In addition to a significant enrichment of mood disorders in this family (~150 family members, corresponding to one third of the pedigree), the pedigree includes individuals with autoimmune thyroid disorder (Jaume et al., 1999) and Ellis-van Creveld Syndrome, an autosomal recessively inherited chondrodysplastic dwarfism (Ginns et al., 2015; McKusick et al., 1964). Our previous work on this pedigree revealed a complex and polygenic inheritance of bipolar disorder with multiple linkage regions and clusters of BD risk-alleles on different haplotypes, supporting a high degree of locus and allelic heterogeneity (Georgi et al., 2014; Kember et al., 2015). Further dissection of the genetic architecture of mental illness in this pedigree has been greatly enhanced by the recently established whole-genome sequence-based imputation reference panel for the Anabaptist population (Based on whole genome sequence for 265 Amish and Mennonite individuals; Anabaptist Genome Reference Panel, AGRP; (Hou et al., 2017)).

In this study, we used the AGRP in combination with genotypes for the ASMAD pedigree (394 family members) to permit the identification of all known disease-causing and loss-of-function (LoF) variants at known Mendelian loci in subjects with mood disorders. In parallel, we performed genomic profiling using polygenic risk scores to establish individuals’ risk for several complex traits and diseases. Long-range phased haplotypes estimated using the extended pedigree permitted the exploration of co-segregation between bipolar risk factors and medical disease loci. We find that a set of 12 Mendelian diseases co-occur in BD individuals more or less frequently than in their unaffected family-members, and that risk scores for metabolic traits are higher in BD individuals, indicating a common genetic etiology for these traits in the extended Amish family.

## Methods

### Sample

The genetic-epidemiologic study of bipolar disorder among the Old Order Amish in Pennsylvania (The Amish Study of Major Affective Disorder) has been well documented (Egeland et al., 1990; Hostetter et al., 1983) and the sample collection methods previously described in detail (Georgi et al., 2014; Kember et al., 2015). Briefly, it consists of a large, extended bipolar disorder pedigree of 700 individuals with a small number of founders. Diagnoses of individuals were made following structured interviews (SADS-L) and a review of medical records by a psychiatric board using strict Research Diagnostic Criteria (RDC) and the Diagnostic and Statistical Manual of Mental Disorders, 4^th^ Edition (DSM-IV) for uniform clinical criteria (Egeland et al., 1990). The majority of affected individuals in the current pedigree are diagnosed as either BPI, BPII, or Major Depressive Disorder. Collection of blood samples followed diagnostic consensus, and lymphoblastoid cell lines were established by the Coriell Institute of Medical Research (CIMR). Signed informed consents were obtained, using language appropriate for Old Order Amish, to a) access medical records for the Amish Study clinicians exclusively to do diagnostic evaluations and clinical studies, and b) to perform collection of blood/tissue samples. In addition, all work contained within this study was approved by the IRB of the Perelman School of Medicine at the University of Pennsylvania.

### Phasing and Imputation

Genotyping was performed on 394 samples from the extended Amish pedigree using Illumina Omni 2.5 M SNP arrays at the Center for Applied Genomics (Children's Hospital of Pennsylvania, Philadelphia, PA). Quality control of the raw genotype calls and imputed genotype calls was conducted using PLINK (Purcell et al., 2007). Individuals were excluded if they (1) had a call rate < 97% or (2) exhibited elevated levels of Mendelian inconsistencies. Variants were excluded from analysis if they: (1) had a call rate < 97% or (2) had minor allele frequencies < 0.002% (i.e., were singletons). Genotypes were phased and imputed with SHAPEIT2 (Delaneau et al., 2011) and IMPUTE2 (Howie et al., 2009), respectively, using the Anabaptist reference panel (Hou et al., 2017). Phasing was performed with SHAPEIT's duoHMM option to account for known familial relationships, using the known ASMAD pedigree, with the genetic map from the HapMap phase II (Frazer et al., 2007), in 5Mb windows. Imputation was performed with IMPUTE2 in 5Mb windows with the following options: - use_prephased_g ‐known_haps_g $SAMPLE.haps ‐phase ‐buffer 500’. Following imputation, genotype dosages were converted to hard calls if above/below a threshold of 1.9/0.1, or otherwise set to missing. In total, 2,379,855 variants were imputed in 394 individuals. Quality control of the imputed calls included removing individuals who had a call rate < 97% (0 individuals), setting Mendelian errors to missing (124,668 variants), and removing variants with a call rate of <99% (120,611 variants) or had minor allele frequencies < 0.002% (869,901 variants). 1,372,783 variants and 394 individuals passed QC. Imputation accuracy was assessed by comparing to whole genome sequence data available for a subset of family members (99.3% concordance).

### Whole genome sequencing

Whole genome sequencing (WGS) for 80 Old Order Amish family members (including 30 parent child trios) was performed by Complete Genomics Inc. (CGI; Mountain View, CA) using a sequence-by-ligation method (Drmanac et al., 2010). Paired-end reads of length 70 bp (35 bp at each end) were mapped to the National Center for Biotechnology Information (NCBI) human reference genome (build 37.2) using a Bayesian mapping pipeline (Carnevali et al., 2012). Variant calls were performed by CGI using version 2.0.3.1 of their pipeline. False discovery rate estimates for SNP calls of the CGI platform are 0.2–0.6% (Drmanac et al., 2010). Gene annotations were based on the NCBI build 37.2 seq_gene file contained in a NCBI annotation build. The variant calls within the WGS were processed using the cgatools software (version 1.5.0, build 31) made available by CGI. The listvar tool was used to generate a master list of the 11.1 M variants present in the 80 Amish samples. The testvar tool was used to determine presence and absence of each variant within the 80 Amish WGS. Only variants with high variant call scores (“VQHIGH” tag in the data files) were included.

### Human disease catalog

The Human Genome Mutation Database (HGMD) catalogs known disease associated variants (http://www.hgmd.org/; (Stenson et al., 2014)). Most of the clinical phenotypes in the database are monogenic diseases. In the June 2013 release it contained 141,000 different variants in ~5,700 genes (“HGMD disease genes”). We examined all variants present in ASMAD in 3456 HGMD disease genes (‘DM’ tag in HGMD).

### Curation of HGMD disease causing variants

251 variants present in ASMAD were annotated as being a 'disease mutation’ in HGMD. In order to further refine this list to identify true disease causing mutations, we first removed all variants found to be present at >1% frequency in 1000 Genomes (Auton et al., 2015) or ExAC (Lek et al., 2016), assuming that disease causing alleles will be rare in a population (total remaining variants=154). Next, we merged this list with annotation from ClinVar (Landrum et al., 2016) and selected all variants annotated as pathogenic (n=62). We then identified whether each variant caused disease in a recessive or dominant model, and used this information to identify individuals in the ASMAD family predicted to display the disease phenotype based on their allelic status (n=25 variants). For these 25 variants, we expanded upon the HGMD and ClinVar annotation by applying criteria recommended by the American College of Medical Genetics and Genomics (ACMG; (Richards et al., 2015)) for the interpretation of sequence variants, and classified the variants into the categories “pathogenic”, “likely pathogenic”, “uncertain significance”, “likely benign” and “benign”. Out of the 25, ACMG criteria classified 3 variants as “benign”, 6 variants as “uncertain significance”, 10 variants as “likely pathogenic” and 6 variants as “pathogenic”. For analyses we removed the 3 variants classified as benign and retained the others (n=22 variants).

### EMMAX and MONSTER

A Balding-Nichols kinship matrix was constructed from the imputed whole genome sequence data following removal of all variants with >5% missing and <1% allele frequency, using the command emmax-kin in the EMMAX package. Association analysis for all variants was performed using EMMAX (Version from February 2012, (Kang et al., 2010)), a statistical test for association analysis using mixed models that accounts for the population structure within the sample. Gene-based association tests were performed using MONSTER (Jiang and McPeek, 2014), a statistical test that generalizes the SKAT-O method and uses a mixed effects model to account for population structure. The kinship matrix used in the EMMAX analysis was also used in the MONSTER analysis.

### Loss of function variants

Variants were annotated using VEP LOFTEE (McLaren et al., 2016). We detect 1177 putative protein truncating (frameshift, splice donor, splice acceptor, and stop-gained) variants. Using filters supplied by LOFTEE, we removed 176 variants (see tables below), resulting in 1001 high confidence loss of function variants (HC-LoF).

| Filter | Description | Number of variants |
| --- | --- | --- |
| END_TRUNC | Variant falls in the last 5% of the transcript | 28 |
| EXON_INTRON_UNDEF | Exon or intron boundaries undefined for this transcript | 1 |
| NON_CAN_SPLICE | Variant falls in non-canonical splice site | 3 |
| NON_CAN_SPLICE_SURR | Variant falls in exon with non-canonical splice site | 7 |

| Flag | Description | Number of variants |
| --- | --- | --- |
| NAGNAG_SITE | Splice acceptor has in-frame AG acceptor site one codon away | 24 |
| SINGLE_EXON | Variant falls in a single exon transcript | 113 |

We identified 167 HC-LoF variants in disease genes, of which 71 are in a homozygous state in one or more individuals. There are 83 rare (<1% in ExAC and 1000G) HC-LoF in disease genes, 17 of which are found in a homozygous state in one or more individuals.

### Association rule discovery

Association rules for the comorbidity of mood disorder and Mendelian diseases in individuals within the ASMAD pedigree were determined using the apriori algorithm from the arules package in R (Hahsler et al., 2005). An itemset was created for each individual, consisting of the affected status for mood disorder (“unaffected” or “broad extended” phenotype) and any comorbid Mendelian disease as determined by the allelic status of the disease causing variant in that individual. Association rules for the frequent itemsets were generated using the apriori command. Rules were then limited to those with either “broad extended” or “unaffected” on the left hand side of the rule (antecedents) and Mendelian diseases on the right hand side of the rule (consequents).

### Polygenic risk scores

A polygenic risk score is generated for each individual as the sum of all variants they carry, weighted by the effect that variant has on phenotype. Polygenic risk scores were generated using the PRSice package (Euesden et al., 2015), with multiple GWAS summary statistics as the base dataset (see Supplemental table 9), and imputed whole genome sequence data in ASMAD as the target dataset. As recommended in the software, we performed p-value informed clumping on the genotype data with an r2 = 0.1 and a distance threshold of 250kb, following exclusion of the MHC region. The optimal p-value threshold for PRS was defined as that which explained the most phenotypic variation for mood disorder in the ASMAD pedigree, out of a set of pre-determined thresholds (p ≤ 0.0001, 0.0005, 0.001, 0.005, 0.01, 0.05, 0.1, 0.5). For traits identified as significantly associated with mood disorder in this pedigree, we show barplots indicating the model fit and p-value for association at each p-value threshold tested (Supplemental figure 29).

### Statistical analysis

Polygenic risk scores for the optimal p-value threshold for each trait were standardized mean=0 and standard deviation=1. Linear mixed model analyses were selected to model outcomes (Narrow phenotype, Broad phenotype, Broad extended phenotype, Depression phenotype) while accounting for relatedness. The analysis was performed using the pedigreemm package in R (Vazquez et al., 2010), with PRS as the independent variable and phenotype as the dependent variable. An empirical kinship matrix, constructed from the genome-wide SNP data, by the Balding-Nicols method using rvtests (Zhan et al., 2016) was fitted as a random effect to account for relatedness between individuals. Risk scores were evaluated for 22 traits, and so a bonferroni corrected p-value<0.0023 (0.05/22) was selected as the level for statistical significance.

### Polygenic transmission disequilibrium

A test for polygenic transmission disequilibrium has recently been described by Weiner et al. (Weiner et al., 2017). We modified this test to allow the comparison of multiple affected and unaffected siblings. We selected all nuclear families with at least one child with Broad phenotype (BPI, BPII, BP:NOS, MDDR, number of families=46). For each family, the polygenic transmission disequilibrium deviation was calculated as follows:

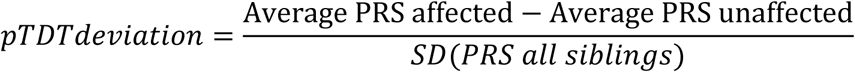

The test statistic for pTDT deviation was then calculated based on all families as follows:

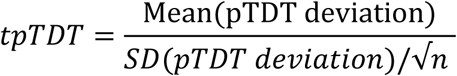

Where n is the number of families.

### Local polygenic risk score

Drawing on work from Shi et al. (Shi et al., 2016), on local genetic correlation, we developed a method for establishing local genetic risk, i.e. genetic risk based on specific regions of the genome. First, we apportioned variants and their corresponding GWAS summary statistics into approximately independent LD blocks (Berisa and Pickrell, 2016). Polygenic risk scores were then generated for each individual in the pedigree based on just the variants within each LD block. For each disease or trait the p-value cut-off for local genetic risk was based on that which explained the most phenotypic variation for mood disorder in the ASMAD pedigree in the genome-wide PRS (calculated as described above). Local risk scores were standardized mean=0 and standard deviation=1. As above, the association of local risk scores with presence of mood disorder were calculated using a linear mixed model with an empirical kinship matrix (constructed from the genome-wide SNP data) fitted as a random effect to account for relatedness between individuals.

## Results

### Identifying the spectrum of mutations in disease genes

By combining SNP genotype data from 394 ASMAD family members with the Anabaptist Genome Reference Panel (Hou et al., 2017) we established high quality phased whole genome sequence for the entire ASMAD extended family. Although available phenotype records for this collection are limited to information about bipolar and related neuropsychiatric disorders (Supplemental table 1), the availability of imputed genotypes allows the identification of ASMAD family members who are carriers of known Mendelian disease variants; and to determine the individual-level burden of common and rare variants in genes associated with a wide range of medical conditions. To ascertain the role of co-morbid diseases in BD in the ASMAD pedigree, we used gene-based and variant-based association tests to explore the role of disease genes in BD enriched loci, identified all variants in known disease genes and investigated segregation of specific disease variants in BD individuals, and used polygenic risk scores to explore the wider genetic architecture of disease in this pedigree.

Disease genes were identified using the Human Genome Mutation Database (HGMD, (Stenson et al., 2014)) by selecting genes with known disease causing mutations (HGMD-DM). The set of HGMD-DM genes were enriched among all genes with predicted damaging variants that are rare in 1000 Genomes and ExAC (<2%) and common in ASMAD (>5%) (57 HGMD-DM out of 239 genes, OR=1.58, p=3.7×10^−3^, two-sided Fisher’s exact test, Supplemental Table 2). Using both gene-based and variant-based association tests we identified loci enriched in affected individuals (none were genome-wide significant, see Supplemental Figures 1 and 2 for Q-Q plots of these results). However, HGMD-DM genes are enriched among both sets of genes (Gene-based test for BD association – MONSTER) and sets of variants (variant-based test for association – EMMAX) that are nominally associated with BD in this pedigree. Out of the top 6 genes that have p-values<0.001, 5 of these are known disease genes (OR=20.3, p=1×10^−3^, two-sided Fisher’s exact test, Supplemental Table 3). Out of the top 44 exonic variants that have p-values <0.001, 17 of these are in known disease genes (OR=2.2, p=1×10^−2^, two-sided Fisher’s exact test, Supplemental Table 4).

In light of these initial results, and the previously noted trend towards a higher burden of CNVs in HGMD-DM genes in subjects with bipolar disorder (Kember et al., 2015), we closely examined variants in disease genes to screen for the presence of Mendelian diseases in the entire pedigree (both affected and unaffected individuals), and to query their contribution to the clinical heterogeneity of BD. Out of 3,456 HGMD-DM genes, 239 contain either known disease causing variants or LoF variants in the ASMAD imputed whole genome sequence. Following extensive curation of variants (see Methods), 62 variants were identified as high confidence disease-causing, of which 17 are known to be more common in the Amish (>5% difference) (Supplemental Figure 3, Supplemental Table 5). Based on the allelic state of these variants in all individuals, we predict that 22 of these are likely to cause disease due to being present in a homozygous (for recessive diseases) or heterozygous (for dominant diseases) state (Supplemental Table 6). Burden of genomically-predicted medical conditions varies between nuclear families, with some families (and individuals within these families) carrying variants for up to 5 Mendelian diseases (Supplemental Figures 4-21). Furthermore, each individual has on average 31 LoF variants in disease genes with 5 disease genes predicted to be completely inactivated.

### BD individuals show comorbidity with a specific set of Mendelian diseases

Overall burden of disease causing variants and LoF variants in disease genes is not associated with BD status in the pedigree. However, the percentage of individuals carrying specific Mendelian disease-causing variants differs between BD individuals and unaffected individuals (Figure 1). To identify if a specific set of Mendelian diseases show comorbidity with BD, we applied association rule discovery to determine diseases that co-occur more frequently than expected with affected or unaffected status of mood disorders (Figure 1, A and B; rule confidence: 0.015-0.123, rule lift: 1.012-1.694). In the ASMAD pedigree, BD individuals are more likely to carry variants causing familial hypercholesterolemia, small fiber neuropathy, and Xerocytosis hereditary (Figure 1, C). Variants causing Adrenocortical hyperplasia and Apolipoprotein C-III deficiency are found more frequently in unaffected individuals.

**Figure 1:**
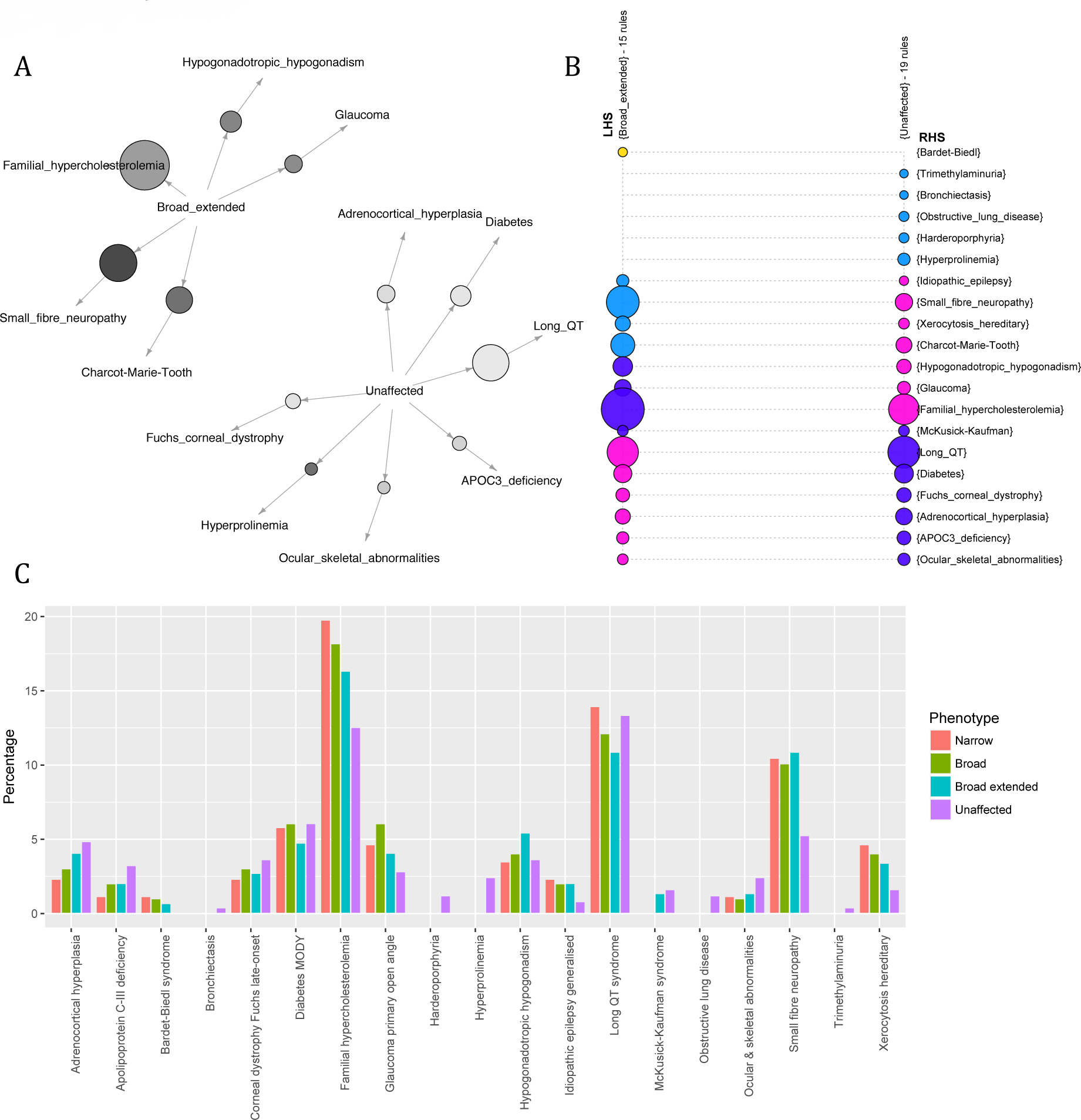
Profile of co-occurring Mendelian diseases varies between affected and unaffected individuals. A and B: Individuals genetically diagnosed with Mendelian diseases. Association rule discovery looks for combinations of variables that occur together more frequently than expected by chance. Diseases that co-occur more than expected differ between individuals affected with mood disorder and unaffected individuals. A: Graph for 12 rules and B: Grouped matrix for 34 rules with mood disorder affected status (either “broad extended” or “unaffected”) as the antecedent and Mendelian disease as the consequent. Size of the circle demonstrates confidence in that rule. Color indicates lift (A: darker=higher lift, B: blue=higher lift, pink=lower lift). C: Individuals carrying disease causing variants. Percentage of Narrow, Broad and Broad extended individuals (see “common polygenic risk for bipolar disorder” in results section for explanation of phenotype) carrying disease causing variants compared to percentage of unaffected individuals with disease causing variants. Some variants are found in a higher percentage of affected individuals.

Half of the Mendelian diseases (6 out of 12) that were identified by association rule discovery as either enriched or depleted in BD individuals are associated with cardiovascular disease or endocrine traits. We therefore explored additional damaging variants associated with lipid phenotypes (Table 1), and found that they are present at a higher frequency in individuals with BD compared to unaffected individuals, including the G574R mutation in *ABCG8* associated with hyperabsorption and sisterolemia, and the R3527Q mutation in *APOB*. In addition to the heterozygotes detected, three homozygotes for the R3527Q mutation were found within the same nuclear family. Strikingly, protective lipid variants are found at a lower frequency in BD individuals compared to unaffected individuals, including a variant in *LPA*, associated with a reduction in thermogenic lipoprotein. Variants associated with heart rhythm (in *KCNH2, SCN5A* and *SNTA1*) show no allele frequency difference between affected and unaffected individuals. We also found that a higher percentage of BD individuals than expected have multiple lipid variants, whereas a lower percentage of unaffected individuals carry multiple lipid variants (Supplemental figure 22). Overall, over one third (34.0%) of affected individuals (broad extended phenotype, dispersed throughout the pedigree) carry at least one variant associated with damaging lipid or iron overload effects, compared to just under one quarter of unaffected individuals (24.7%).

**Table 1:**
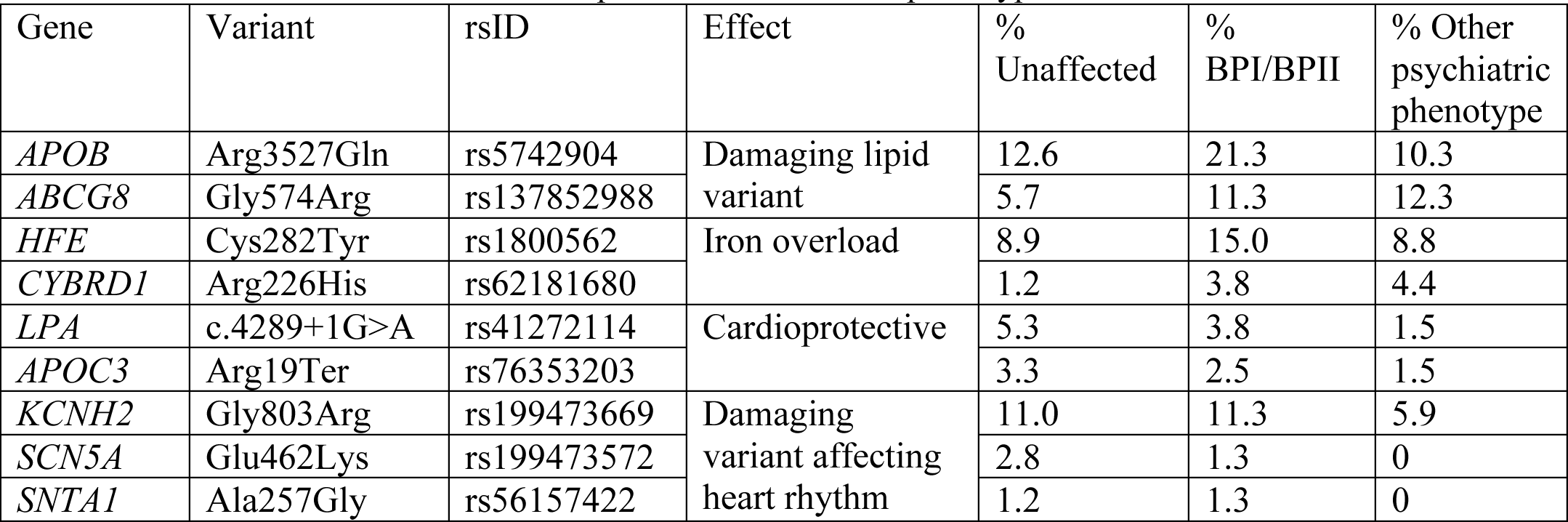
Variants associated with lipid and cardiovascular phenotypes

### Common polygenic risk for bipolar disorder

To compare the genetic architecture of BD in the Amish with a non-founder population, and to further explore the enrichment of variants associated with lipid traits in BD subjects, we sought to measure the aggregate effect of common variants underlying both psychiatric and endocrine/metabolic traits in family members with and without mood disorders. To achieve this, we evaluated different phenotypic models from the most defined (BPI individuals only) to the most general (all individuals with any psychiatric phenotype, including minor psychiatric symptoms). Heritability estimates generated for this pedigree previously have shown significant heritability for mood disorder phenotypes (Georgi et al., 2014). Using a refined pedigree structure, updated from accurate genealogical records, we estimated narrow sense heritability for the following phenotype models: narrow (BPI and BPII only), broad (BPI, BPII, and MDDR), broad extended (presence of any psychiatric phenotype), depression (MDDR, MDD, and Minor Depression), and well. We found significant heritability for all phenotype models (Supplemental Table 7), with broad extended being the most heritable phenotype (h^2^=0.81, p=9.76×10^−10^). We generated polygenic risk scores based on GWAS summary data (from Europeans) for multiple traits (see Supplemental Table 9 for full list) and tested for an association between the risk score and presence of mood disorder (Figure 2, Table 2, Supplemental Table 8). Polygenic risk scores (PRS) incorporate multiple SNPs, including those that do not reach the threshold for genome-wide significance, to predict an individual's risk for disorder. First, we used summary statistics from a GWAS for bipolar disorder by the Psychiatric Genomics Consortium (PGC BD2, (Stahl et al., 2017)), to generate risk scores for bipolar disorder for all individuals in the ASMAD pedigree. Linear mixed model analyses (Vazquez et al., 2010) permitted an association test for risk scores while accounting for relatedness within the pedigree. We tested the difference in risk scores between unaffected and affected individuals for the above phenotype models.

**Figure 2:**
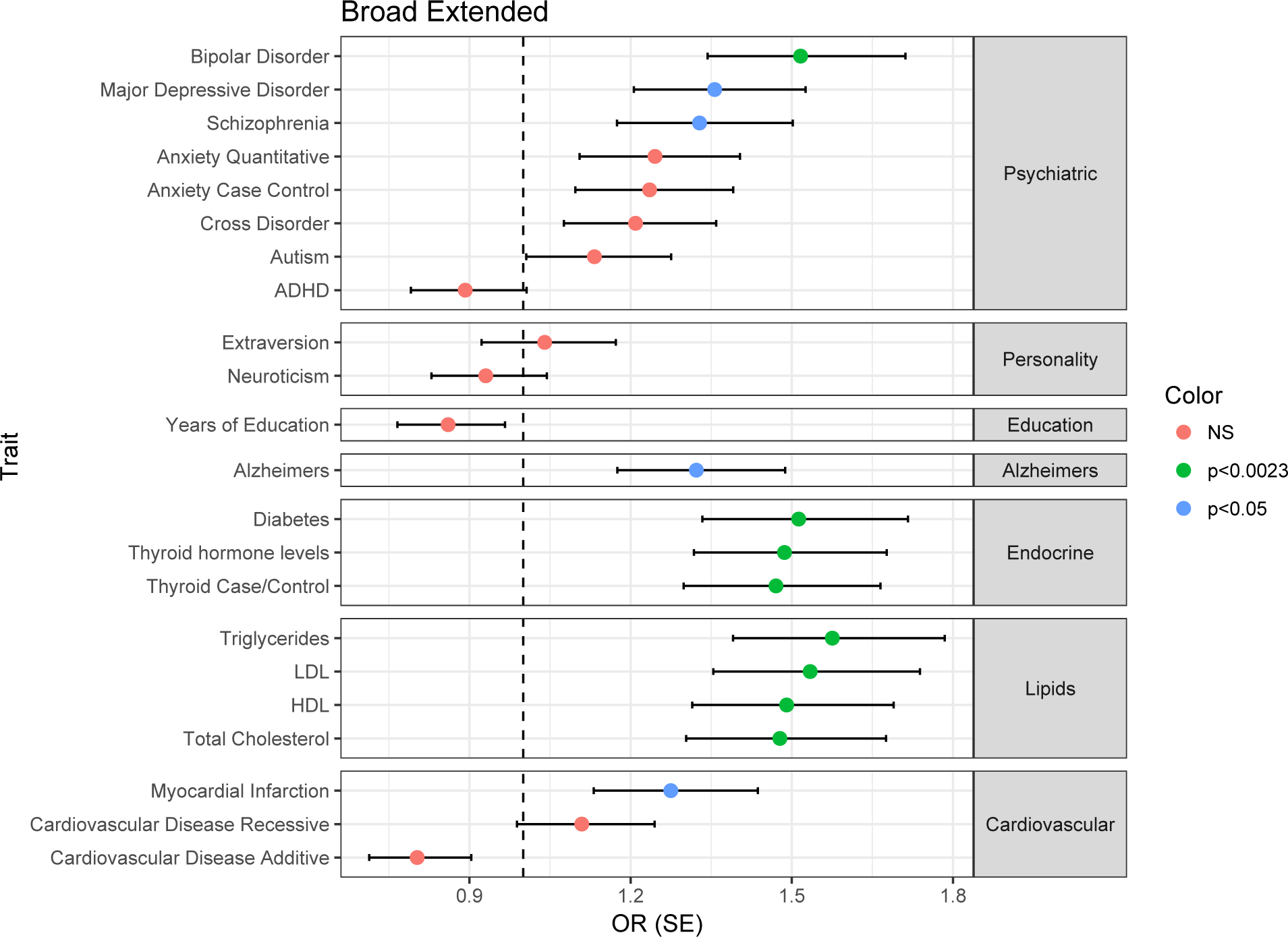
Association of polygenic risk scores for multiple traits with bipolar disorder (broad extended phenotype) in ASMAD. Risk scores for all traits tested are displayed, split by trait type. Circles indicate the association of the risk scores with mood disorder in the ASMAD pedigree, color of circles shows the level of significance, and error bars show +/− standard error.

**Table 2:** The associations between polygenic risk scores for the trait and mood disorder (broad extended phenotype) with the largest effect size are presented. Statistically significant p-values following bonferroni correction (p<0.0023) are shown in bold.

| Category | Trait/GWAS | Best fit threshold | # SNPs | R<sup>2</sup> | P-value | Beta adjusting for relatedness | P-value adjusting for relatedness |
| --- | --- | --- | --- | --- | --- | --- | --- |
| Psychiatric Disorder | **Bipolar Disorder (PGC BD2)** | **0.05** | **87318** | **0.04900** | **0.00021** | **0.416** | **0.00059** |
|  | Major Depression (PGC MDD) | 0.1 | 140450 | 0.02865 | 0.00427 | 0.305 | 0.00945 |
|  | Schizophrenia (PGC SCZ2) | 0.5 | 453646 | 0.01937 | 0.01824 | 0.284 | 0.02079 |
|  | Autism (PGC AUT) | 0.001 | 1284 | 0.00614 | 0.18422 | 0.125 | 0.29455 |
|  | Cross disorder (PGC) | 0.001 | 799 | 0.01063 | 0.08162 | 0.190 | 0.10413 |
|  | Anxiety (ANGST Case-control) | 0.005 | 4722 | 0.01488 | 0.03899 | 0.211 | 0.07468 |
|  | Anxiety (ANGST Quantitative phenotype) | 0.5 | 149313 | 0.01437 | 0.04375 | 0.219 | 0.06626 |
|  | ADHD (EAGLE) | 0.0005 | 383 | 0.00638 | 0.17754 | -0.114 | 0.34360 |
| Personality | Neuroticism (GPC2) | 0.05 | 36987 | 0.00344 | 0.31990 | -0.072 | 0.53126 |
|  | Extraversion (GPC2) | 0.5 | 233177 | 0.00073 | 0.64621 | 0.039 | 0.74262 |
| Educational attainment | SSGAC (EA 2016) | 0.5 | 442230 | 0.01191 | 0.06386 | -0.150 | 0.19475 |
| Alzheimer's | IGAP Stage 1 | 0.1 | 130780 | 0.02542 | 0.00708 | 0.279 | 0.01770 |
| Diabetes | **GoT2D (DIAGRAM)** | **0.5** | **98118** | **0.04620** | **0.00033** | **0.414** | **0.00103** |
| Thyroid | **Thyroid hormone levels** | **0.0001** | **118** | **0.04207** | **0.00085** | **0.386** | **0.00191** |
|  | **Thyroid antibodies** | **0.001** | **827** | **0.04986** | **0.00025** | **0.396** | **0.00010** |
| Lipids | **HDL (Global Lipids)** | **0.5** | **75151** | **0.04369** | **0.00046** | **0.399** | **0.00145** |
|  | **LDL (Global Lipids)** | **0.05** | **12403** | **0.04911** | **0.00020** | **0.428** | **0.00062** |
|  | **TC (Global Lipids)** | **0.5** | **77622** | **0.04227** | **0.00056** | **0.391** | **0.00187** |
|  | **TG (Global Lipids)** | **0.1** | **21255** | **0.05670** | **0.00007** | **0.455** | **0.00027** |
| Cardiovascular Disease | CAD additive | 0.0001 | 121 | 0.01618 | 0.03151 | -0.220 | 0.06265 |
|  | CAD recessive | 0.001 | 1428 | 0.00871 | 0.11340 | 0.104 | 0.36892 |
|  | MI additive | 0.001 | 2403 | 0.01154 | 0.06913 | 0.243 | 0.04220 |

Bipolar disorder risk score was significantly associated with affected phenotype for broad extended (β=0.416, SE=0.121, p=5.9×10^−4^), broad (β=0.523, SE=0.129, p=4.8×10^−5^), and narrow phenotypes (β=0.499, SE=0.138, p=3×10^−4^), but not for depression phenotype (β=0.273, SE=0.155, p=7.8×10^−2^). To verify this result, we used a modified polygenic transmission disequilibrium test (see Methods) and found a statistically significant deviation in risk scores, with affected individuals having a higher polygenic risk than their unaffected siblings (mean deviation 0.27, p=3×10^−2^). This increase is mostly driven by higher risk scores of individuals with BPI, BPII, BP:NOS, and MDDR (Supplemental Figure 23). The percentage of affected individuals, on average, increases with increasing deciles of PRS (Figure 3A). Furthermore, the percentage of individuals with BPI, BPII and BP:NOS is highest in the 10^th^ PRS decile, whereas the percentage of individuals with minor depression or other non-specified disorder is lowest in this decile (Figure 3B). Offspring with an affected parent (either one or both parents affected) have higher risk scores than offspring from unaffected parents (Supplemental Figure 24). Furthermore, descendants that can trace their lineage back to the two pioneer members of this pedigree (“in family”) have significantly higher risk scores for bipolar disorder than Amish individuals who are “married in” (β=1.653, SE=0.370, p=8×10^−6^, Supplemental Figure 25), suggesting that the pioneer individuals carried high risk for bipolar disorder. Risk scores vary across the pedigree, although they are more similar within nuclear families (Supplemental Figure 26).

**Figure 3:**
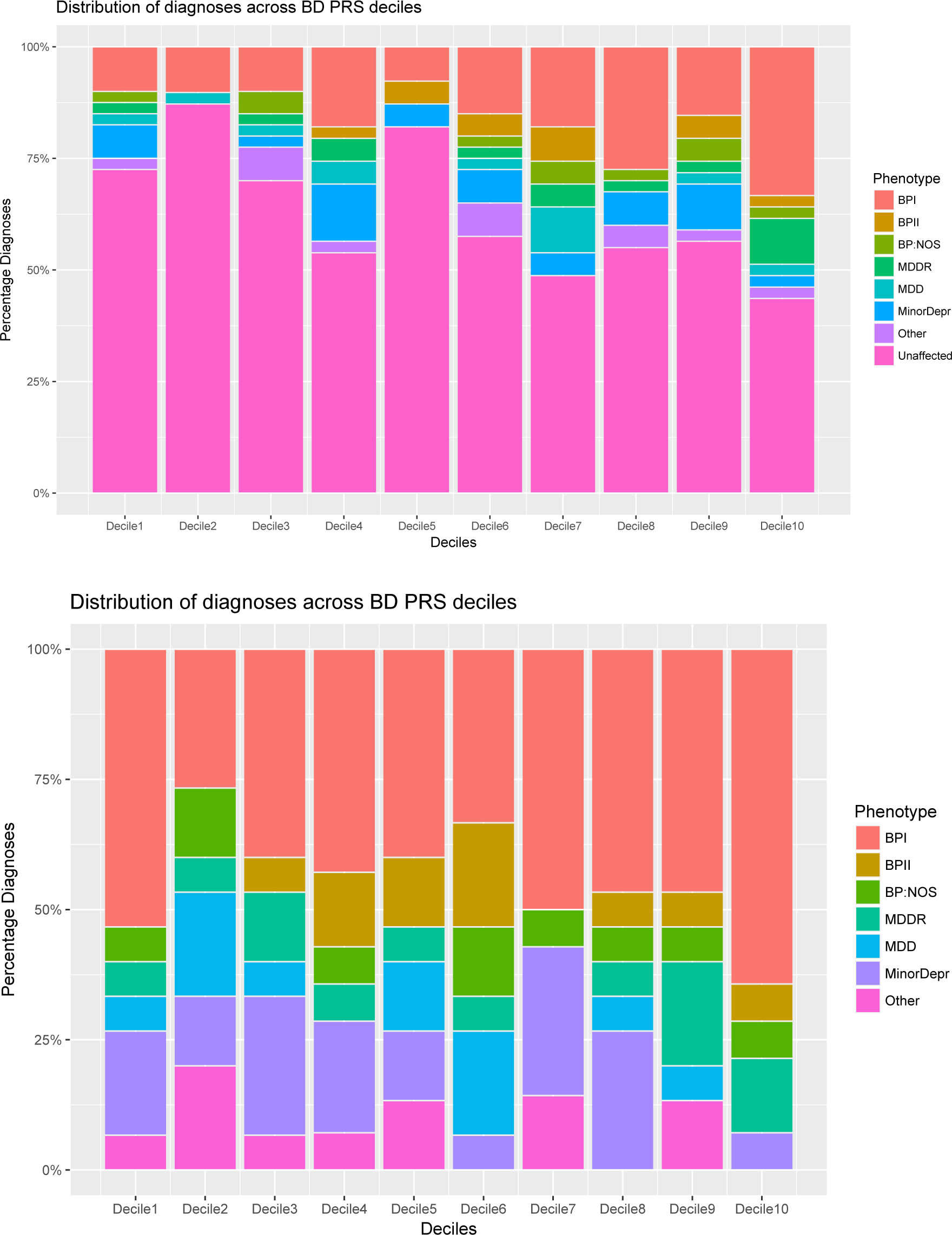
Percentage of individuals with affected status for each decile of polygenic risk score. A: All individuals, including unaffected individuals, are shown. B: Affected individuals only are shown.

### Common risk variants for disease suggest pleiotropy between common complex traits and bipolar disorder

Risk score analyses can also be used to query genetic associations between complex disorders that have shown comorbidity in individuals but were otherwise thought to be unrelated (Wray et al., 2014). Therefore we compiled published GWAS summary statistics data (Supplemental table 9) and generated risk scores for 22 additional traits, including common psychiatric disorders, personality measures, educational attainment, Alzheimer's, Type 2 diabetes, autoimmune thyroid disease, lipid traits, and cardiovascular disease (Figure 2, Table 2). Risk scores for clinical autoimmune thyroid disease, lipid traits (Total cholesterol, Triglycerides, LDL, HDL) and type 2 diabetes were found to be significantly higher in individuals with mood disorder even after accounting for pedigree structure, suggesting that in this pedigree there are shared loci between bipolar disorder and both lipid traits and diabetes. Thyroid hormone levels, positive thyroid antibodies, HDL, LDL, triglyceride, total cholesterol, and diabetes risk scores were significantly associated with affected phenotype, but only thyroid hormone, LDL and triglyceride risk scores were nominally associated with depression phenotype (for full results, see Supplemental table 8).

In the extended Amish pedigree, risk scores for bipolar disorder are positively correlated with risk scores for lipid traits (TG r=0.55, p<2.2×10^−16^, TC r=0.55, p<2.2×10^−16^, LDL r=0.52, p<2.2×10^−16^, HDL r=0.57, p<2.2×10^−16^, Supplemental Figure 27), diabetes (r=0.54, p<2.2×10^−16^), thyroid levels (thyroid hormone levels r=0.31, p=1.6×10^−10^), major depression (r=0.39, p=6.4×10^−16^), schizophrenia (r=0.55, p<2.2×10^−16^), and Alzheimer's disease (r=0.40, p=2.9×10^−16^), and negatively correlated with years of education (r=-0.21, p=2×10^−5^), suggesting an overlap between loci that contribute to risk for these traits. Furthermore, we explored the effect of inbreeding on risk score values, as increased homozygosity in offspring of closely related parents is known to increase the incidence of recessive Mendelian disorders, but the effect on complex disorders is less well quantified. We found that higher inbreeding values (measured as a higher number of observed homozygous genotypes than expected), but not increased average length of homozygous regions, are correlated with an increased risk score for bipolar disorder (r=0.69, p=2.2×10^−15^, Supplemental Figure 28), suggesting a role for increased homozygosity in common disease risk.

### Identifying pleiotropic regions of the genome

To establish which genomic regions contribute to the genome-wide risk score differences between affected and unaffected individuals, we calculated risk scores for each approximately independent LD block (n=1,703) as defined by Berisa and Pickrell (2016). This allowed us to partition the genome and analyze risk for each local region to identify whether a) a single region or multiple regions are underlying the overall difference in polygenic risk between affected and unaffected family members, and b) if the regions underlying differences in risk are the same between the different traits and disorders analyzed. We calculated local polygenic risk for bipolar disorder, total cholesterol, LDL, HDL, triglycerides, diabetes, and thyroid disorder, and tested for association with mood disorder (Broad extended phenotype) using linear mixed model analyses to account for relatedness (as above).

We identified a single genomic region on chromosome 15 for bipolar disorder local polygenic risk (15q25.2-q25.3) that was significantly associated with mood disorder in the ASMAD pedigree (β=0.495, SE=0.113, p=1.19×10^−5^) following bonferroni correction to account for the number of regions tested. This region contains one of the genome wide significant loci identified in the PGC BD2 study (Stahl et al., 2017), near *ZNF592*, a gene thought to play a role in cerebellar development. None of the other regions for any of the risk scores were significant following correction for multiple testing. We therefore explored all regions reaching a threshold of p<0.01, to discover if any of the nominally significant regions overlapped between traits and diseases. Using this threshold we identified 19 genomic regions for bipolar disorder, 24 for total cholesterol, 25 for LDL, 25 for HDL, 21 for triglycerides, 25 for diabetes, 1 for presence of thyroid antibodies, and 4 for thyroid hormone levels (Figure 4, Supplemental Table 10). Of the 144 regions identified, there were 121 unique regions, with 17 regions overlapping between traits.

**Figure 4:**
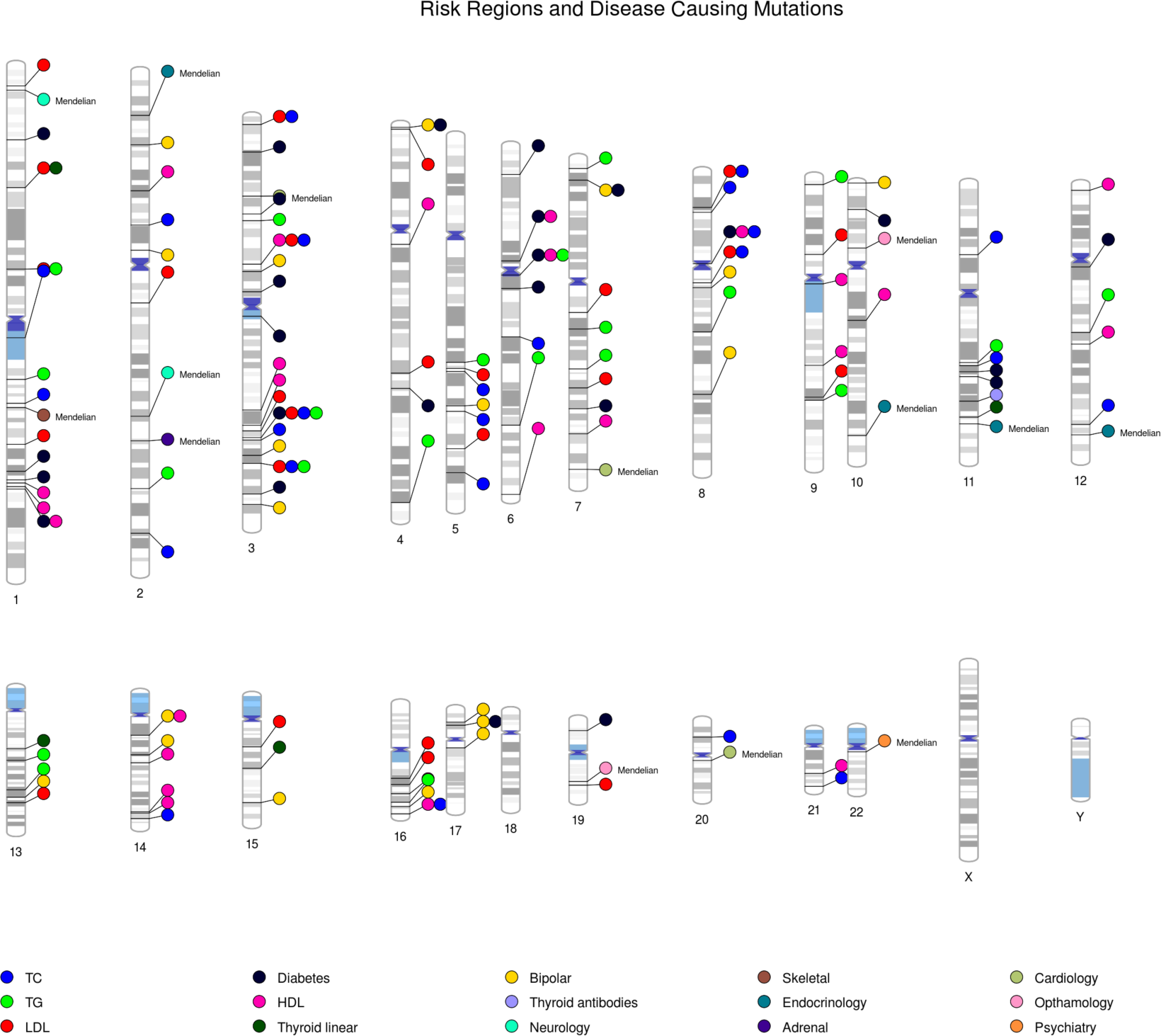
Locations of genomic regions that contribute to the genome-wide risk score differences between affected and unaffected individuals for bipolar disorder (yellow), diabetes (black), HDL (pink), LDL (red), Total cholesterol (blue), triglycerides (light green) thyroid antibodies (lavender) and thyroid hormone (dark green). The location of Mendelian variants which are associated with either affected or unaffected status in this pedigree are also included (denoted by the notation: Mendelian).

The telomeric region on chromosome 4 (4p16.4) harbored local risk for both bipolar disorder and diabetes that associated with mood disorder status in the pedigree (BD: β=0.449, SE=0.118, p=1.4×10^−4^, diabetes: β=0.323, SE=0.114, p=4.6×10^−3^). This region was previously identified as the region with the highest linkage with bipolar disorder in this family (LOD 3.95) (Georgi et al., 2014). Bipolar disorder and diabetes local polygenic risk regions also overlapped on chromosome 7 (7p21.3), a region that contains the gene *NDUFA4* and has previously been associated with a psychosis phenotype (Bramon et al., 2014), and chromosome 17 (17p12), a region containing the genes *MYOCD*, *ARHGAP44*, and *ELAC2,* with no previous associations to mood disorders. Bipolar disorder and HDL local polygenic risk associated with mood disorder overlapped on chromosome 14 (14q12-13.1), a region harboring the gene *NPAS3*, which has previously been associated with schizophrenia susceptibility in linkage (Chiu et al., 2002) and association studies (Huang et al., 2010; Pickard et al., 2009). Diabetes, HDL, LDL, triglycerides, total cholesterol, and thyroid disorder local polygenic risk regions associated with mood disorder in this pedigree overlapped at multiple locations that were not additionally associated with polygenic bipolar local risk scores. Finally, the other regions were not found to overlap between traits, suggesting that the association of bipolar disorder with risk for other traits in this pedigree is due to multiple pleiotropic regions spread across the genome.

## Discussion

Comorbidity of medical disease in individuals with psychiatric disorder is a major contributor to early mortality and severity of phenotype. Although previously attributed to poor lifestyle choices, recent genetic evidence suggests that pleiotropic loci that predispose to both mental illness and non-psychiatric disease may underlie at least some comorbidities. Here we present a large multigenerational pedigree significantly enriched for Bipolar and related mood disorders, that we further genetically diagnose with segregating Mendelian disease variation. In addition to identifying a unique pattern of co-segregating low-frequency Mendelian disease variants associated with mood disorder status, polygenic risk profiling, using several sets of common variants associated with a wide range of complex diseases and traits, provides strong support for a shared genetic architecture between mood disorders and specific metabolic and endocrine traits. Although our detected associations may be unique to the Amish extended pedigree, we propose that such patterns of phenotypic and genetically-defined pleiotropy may enable subtyping of complex diseases and facilitate their genetic dissection.

The expectation in studies of complex disease in founder populations is that low-frequency disease-causing variants with strong effect sizes, that are rare in the general population but enriched or specific to the isolate, can be more easily identified compared to studies in outbred populations. However, previously published findings from the ASMAD population (Georgi et al., 2014; Ginns et al., 2015; Kember et al., 2015) reveal a complex polygenic mode of inheritance, in line with findings from general population studies. Here we corroborate this conclusion by reporting that bipolar disorder polygenic risk profiling unequivocally supports a shared genetic architecture (i.e. shared common risk variants) between bipolar disorder in the Amish and in an outbred population of European ancestry. Furthermore, as expected from studies on the shared genetic etiology of psychiatric disorders (Lee et al., 2013), polygenic risk scores for major depressive disorder are also associated with bipolar disorder in this pedigree. Interestingly, despite the high genetic correlation between bipolar disorder and schizophrenia in the general population, the schizophrenia polygenic risk score was limited in its predictive ability in the ASMAD sample (R^2^=0.019). This may reflect an ascertainment bias in recruitment or a unique aspect of the phenotype in this extended pedigree. For example, out of 195 individuals with any reported psychiatric condition in the family, only a single individual presented with schizoaffective disorder and none with schizophrenia. Similarly, a study of major psychiatric disorders in mid-Western Amish in Ohio and Indiana reported lower rates of psychosis compared to other populations (Hou et al., 2013).

Our previous analysis of regions of homozygosity did not reveal recessive risk variants (SNV or CNVs) for BD in this pedigree (Georgi et al., 2014; Kember et al., 2015). Conversely, here we report that polygenic risk scores for bipolar disorder and several metabolic traits are positively correlated with the inbreeding co-efficient. Interestingly, in an effort to investigate genetic diversity within Anabaptist groups represented in the Anabaptist Genome Reference Panel (AGRP) (Hou et al., 2017), we showed that Amish living in Lancaster County, Pennsylvania, where the ASMAD pedigree originates, have the longest homozygous-by-descent-regions. Therefore, we suggest that reduced genetic diversity may underlie the accumulation of both common and low-frequency, homozygous and compound heterozygous, risk alleles for a range of diseases and traits.

Our identification of individuals who should present with a Mendelian disease, based on their carrier status of highly penetrant Mendelian variants, established a set of diseases that differ in frequency between those with and without mood disorder. Moreover, individuals carrying these variants in a manner insufficient for presentation of disease also demonstrated differences in mood disorder status. For example, a dominant familial hypercholesterolemia-associated APOB R3527Q variant, which has been previously identified as being carried by around 12% of Old Order Amish individuals (Shen et al., 2010), was found in 19.7% of individuals with BPI, BPII or BP:NOS. This finding suggests that rather than one trait having a causal effect on the other (e.g. hypercholesterolemia “causing” bipolar disorder) it is more likely that a gene variant, or the haplotype containing this Mendelian variant harbors other co-segregating variants, that convey risk for both disorders.

To further explore the biological implications of our finding that a specific set of Mendelian variants are enriched in individuals with mood disorders, we generated risk scores for multiple traits using common disease associated variants. We identified a number of complex metabolic and endocrinological diseases (lipid traits, diabetes and clinical thyroid disease) that were significantly associated with mood disorder in this pedigree. The relationship between thyroid disorders and mood disorders has been acknowledged for many years (Whybrow et al., 1969), with thyroid function associated with depressive symptoms, anxiety, and mania (Chakrabarti, 2011). While metabolic syndrome in general (Fiedorowicz et al., 2008; Vancampfort et al., 2013), and diabetes (Golden et al., 2008) and lipid levels (Enger et al., 2013; Patel et al., 2007) specifically, have been shown to be elevated in individuals with mood disorder, cross-trait analysis has not provided evidence for a significant genetic correlation between bipolar disorder and any metabolic trait (Bulik-Sullivan et al., 2015). Our study on individual level data in a large extended family reveals evidence for a genome-wide genetic correlation between mood disorder and specific metabolic traits. As this finding has not yet been replicated in an outbred population, we hypothesize that BD in the ASMAD family could represent a subtype of BD with high levels of metabolic and endocrinological disorders. This has been termed subgroup heterogeneity (Han et al., 2016), where a genetically distinct subset of individuals within a patient cohort is also genetically similar to individuals with another disease. There is already emerging evidence that such a subgroup exists for major depressive disorder (Howard et al., 2017), and the identification of a similar subgroup in BD cohorts may help explain the epidemiological findings of associations between these disorders.

In an effort to quantify the regions which contribute to differences in risk scores, we take advantage of the long haplotypes shared between multiple related individuals to analyze risk scores for each approximately independent LD block across each chromosome and identify regions for which risk scores differ between individuals with mood disorder and their unaffected relatives. We identified a relatively small number of regions for each trait, covering 0.05-2.35% of the genome, that are associated with differences in risk scores at a p-value<0.01. While 17 regions overlap between traits at this nominal cutoff, the majority of these regions are unique to each trait; out of the 19 regions identified as driving bipolar risk differences between affected and unaffected individuals, only four of these overlap with another trait. This suggests that, overall; regions that underlie the difference between affected and unaffected individuals polygenic risk for bipolar disorder are not the same regions that underlie the difference between affected and unaffected individuals polygenic risk for metabolic traits. However, we assume that each of these regions are pleiotropic for both the trait that the risk score has been created for and for mood disorder phenotype in this pedigree, as the regions are carrying risk variants for the trait tested based on summary statistics from an independent data set, and the risk scores are different between affected and unaffected individuals in this pedigree. Using this method we are unable to determine if the variants associated with the metabolic or endocrine trait are also influencing risk for bipolar disorder in this family (pleiotropy at the level of the variant), or if the region contains haplotypes carrying both variants that influence risk for the trait in addition to variants that influence risk for mood disorder (pleiotropy at the level of the haplotype).

We acknowledge several limitations within our current work that highlight avenues for future study. There are many cases of disease genes with different alleles leading to different phenotypes or a specific allele leading to a range of diverse phenotypes (Lupski et al., 2011). Although there are clear examples of homozygous variants causing a recessive Mendelian disease and a complex condition in carriers (Lupski et al., 2011), our study does not provide the resolution required to distinguish between a single gene vs. haplotype effect. Specifically, we can't determine whether variation in the Mendelian disease gene alone, or variation at the level of the haplotype, is responsible for the shared genetic etiology of disease in this population. We expect that the long shared haplotypes observed in a genetic isolate are more likely to harbor multiple disease-contributing variants, and therefore produce the observed genetic pleiotropy. Furthermore, multiple mechanisms, such as zygosity, epistasis and other interacting genes (genetic background) or environmental effects may modify expressivity or penetrance of a specific allele (Zhu et al., 2014). While every attempt was made to limit our analyses to Mendelian variants predicted to cause disease, the presentation of Mendelian disease in members of the ASMAD pedigree could not be confirmed due to restrictions on re-contacting individuals in this legacy collection. As pleiotropic alleles continue to be identified, future studies would benefit from broadly phenotyping cases to fully capture the combination of traits and diseases present in each individual.

In conclusion, we demonstrate a case of genetic pleiotropy between a complex psychiatric disease with both Mendelian and complex metabolic and endocrine traits. We suggest this indicates a common genetic etiology for these traits in the extended Amish family. While our specific findings may not be extendable to other populations, we propose that each pedigree or population segregating psychiatric disorders will have unique combinations of additional medical traits. Taken together, our results denote that medical comorbidity between complex diseases and Mendelian disorders arises as a combination of chromosomal proximity of disease causing variants and pleiotropy of disease genes. Elucidating these patterns could enhance the ability to identify regions and variants that contribute to disease in each unique population.

## Acknowledgments

The authors are indebted to the members of the Old Order Amish settlements who participated in the Amish Study of Major Affective Disorder and Dr. Janice Egeland who designed and directed the initial recruitments and clinical evaluation. This study was supported by the NIH grant R01MH093415, and supported in part by the NIMH Intramural Research Program. The authors are thankful to Drs. Wade Berrettini, Elliot Gershon, Danish Saleheen, Iain Mathieson, Bogdan Pasaniuc and Daniel Rader for informative discussions and their feedback on the manuscript. We are grateful to Marco Medici and Anne Cappola for the summary statistics for thyroid traits, the Psychiatric Genomics Consortium (in particular the Bipolar Disorder, MDD, and SCZ subgroups) and the many other studies and consortia that made their summary statistics readily available.

